# Engineering living immunotherapeutic agents for improved cancer treatment

**DOI:** 10.1101/2023.03.31.535049

**Authors:** Tinotenda Gwisai, Sina Günther, Matej Vizovisek, Mira Jacobs, Simone Schuerle

## Abstract

Bacteria-based biohybrid agents are emerging as a promising strategy for cancer therapy due to their ability to actively target tumors, trigger localized inflammation and induce tumor regression. There has been growing interest in using bacteria that are responsive to external cues, such as magnetic fields, to facilitate the formation of robust colonies in tumor the achieve the threshold for clinical efficacy. Several studied have demonstrated the potential of innately magnetically responsive bacteria, known as magnetotactic bacteria (MTB), as steerable agents, however, their immunostimulatory properties and therapeutic effects are yet to be explored. Here, we characterize key properties of human immune cell responses and the behavior of the MTB strain *Magnetospirillum magneticum* AMB-1 in physiological environments. This work investigates the ability of MTB to maintain magnetic properties, viability in whole blood, cytokine production by macrophages, and the ability to increase uptake of cancer cell material by dendritic cells. This study also explores the use of MTB-liposome complexes for effective delivery of therapeutic payloads. Overall, this study establishes the potential of MTB as a versatile, combined drug delivery platform for immune-mediated cancer therapy.

## Introduction

Cell-based therapies are ushering in a new era of cancer treatment in which the innate tumor-targeting attributes of living agents are leveraged to overcome current limitations of conventional treatment (*1*). Accumulation of systemically administered therapeutics in tumors is currently hindered by diffusion-limited transport and biological barriers, including highly irregular vasculature and elevated interstitial fluid pressure, both of which impede drug penetration (*2–4*). In contrast, biohybrid microrobots, that combine living agents with synthetic materials, have the ability to actively target tumors in response to cues stemming from the unique chemical composition of tumors and navigate through various tissues using self-propulsion (*5*).

Bacteria-based biohybrid systems have reemerged as particularly promising cancer therapy agents due to their ability to selectively colonize tumors, the intrinsic cytotoxicity of certain strains and their capacity to evoke an immune-mediated antitumor response (*6*). Since tumors can develop methods to evade immune recognition, including the production of immunosuppressive cytokines (*7*), bacterial agents can be used to overcome some of these mechanisms. Pathogen associated molecular patterns on bacteria, such as lipopolysaccharides (LPS), are recognized by innate immune cells and this can induce the production of various proinflammatory cytokines (*8*). The effects of bacteria-based cancer therapy can be complemented or enhanced by payload delivery. Recently, non-pathogenic *E. coli* Nissle 1917 (EcN) has been engineered to produce stimulator of interferon genes (STING) agonists (*9*), as well as programmed cell death–ligand 1 (PD-L1), cytotoxic T lymphocyte–associated protein-4 (CTLA-4) and CD47 (*10, 11*) nanobody antagonists to trigger localized inflammation and induce tumor regression.

While preclinical studies have shown that bacteria-based microrobots can be used to trigger effective tumor elimination, suboptimal clinical responses have hampered the translation of this approach. Since sufficient tumor colonization is a prerequisite for successful therapy, interest has grown in using external cues to aid the formation of robust colonies (*12, 13*). Responsiveness to external stimuli gives dual-targeting functionality to bacterial agents that are equipped with intrinsic tumor homing mechanisms. Magnetotactic bacteria (MTB), a group of bacteria that biomineralize magnetic nanoparticles arranged in chains, have driven interest in using externally applied magnetic fields to control bacteria. The MTB strain MC-1 was found to preferentially accumulate in hypoxic tumor regions following peritumoral injection and guidance using externally applied DMF (*14*). The MTB strain AMB-1 has also been used as a magnetically controllable motile carrier of indocyanine green nanoparticles for photothermal therapy (*13*). Recent studies have also explored the control of genetically engineered EcN that have been functionalized with magnetic nanoparticles for photothermal therapy and magnetothermal tumor ablation (*15, 16*). Unlike other strains of bacteria, MTB uniquely combine tumor homing capacity with intrinsic production of anisotropic magnetic chains, making them suitable for manipulation with a range of magnetic stimuli, including rotating magnetic fields. For instance, our recent work demonstrated the efficacy of a hybrid control strategy that combines magnetic torque-driven motion via rotating magnetic fields followed by taxis-based navigation to significantly enhance tumor infiltration of MTB *in vivo* (*17*). Despite the great promise of MTB as bacterial microrobots, illustrated by numerous previous studies, interactions between host immunity and MTB are yet to be characterized and their potential as potent, living anticancer agents is yet to be established.

To employ MTB for cancer therapy, it is imperative to characterize their immunostimulatory properties and behavior in physiological environments. In this work, we first investigated the ability of the MTB strain *Magnetospirillum magneticum* AMB-1 to proliferate and maintain their magnetic properties under physiological conditions, as well as their viability in whole blood. We then examined selected key properties of human innate immune cells in response to stimulation with MTB. Since cytokine production is essential for mounting effective immune-mediated tumor clearance, the cytokines secreted by THP-1 derived macrophages stimulated with MTB were evaluated. Next, we studied the ability of MTB to increase the uptake of cancer cell material by monocyte-derived dendritic cells (moDCs), as these cells play a key role in triggering an adaptive immune response. Lastly, the use of MTB-liposomes complexes (MTB-LP) for effective delivery of therapeutic payloads was investigated. Overall, this study establishes the potential of the MTB-LP system as a versatile, combined drug delivery platform and lays the foundation for further characterization of the efficacy of MTB for immune-mediated cancer therapy.

## Results and Discussion

### MTB proliferation under physiological conditions and clearance from whole blood

Since MTB are aquatic bacteria and an atypical strain for bacterial cancer therapy, we first investigated the ability of MTB to proliferate and maintain their magnetic properties under physiological conditions. In addition to standard MTB culture conditions of 30 °C in magnetic spirillum growth medium (MSGM), other conditions were investigated where the type of medium or the temperature was modified (Figure S1). Optical density was measured over 10 days of incubation and the corresponding bacterial concentrations were computed. The typical characteristics of MTB proliferation, where exponential growth in the first 2 days was followed by a stationary phase, were observed in all conditions (Figure 2A). Samples cultured at 37 °C had higher MTB concentrations than cultures at 30 °C, and samples cultured in DMEM had higher concentrations than cultures in MSGM.

**Figure 1:**
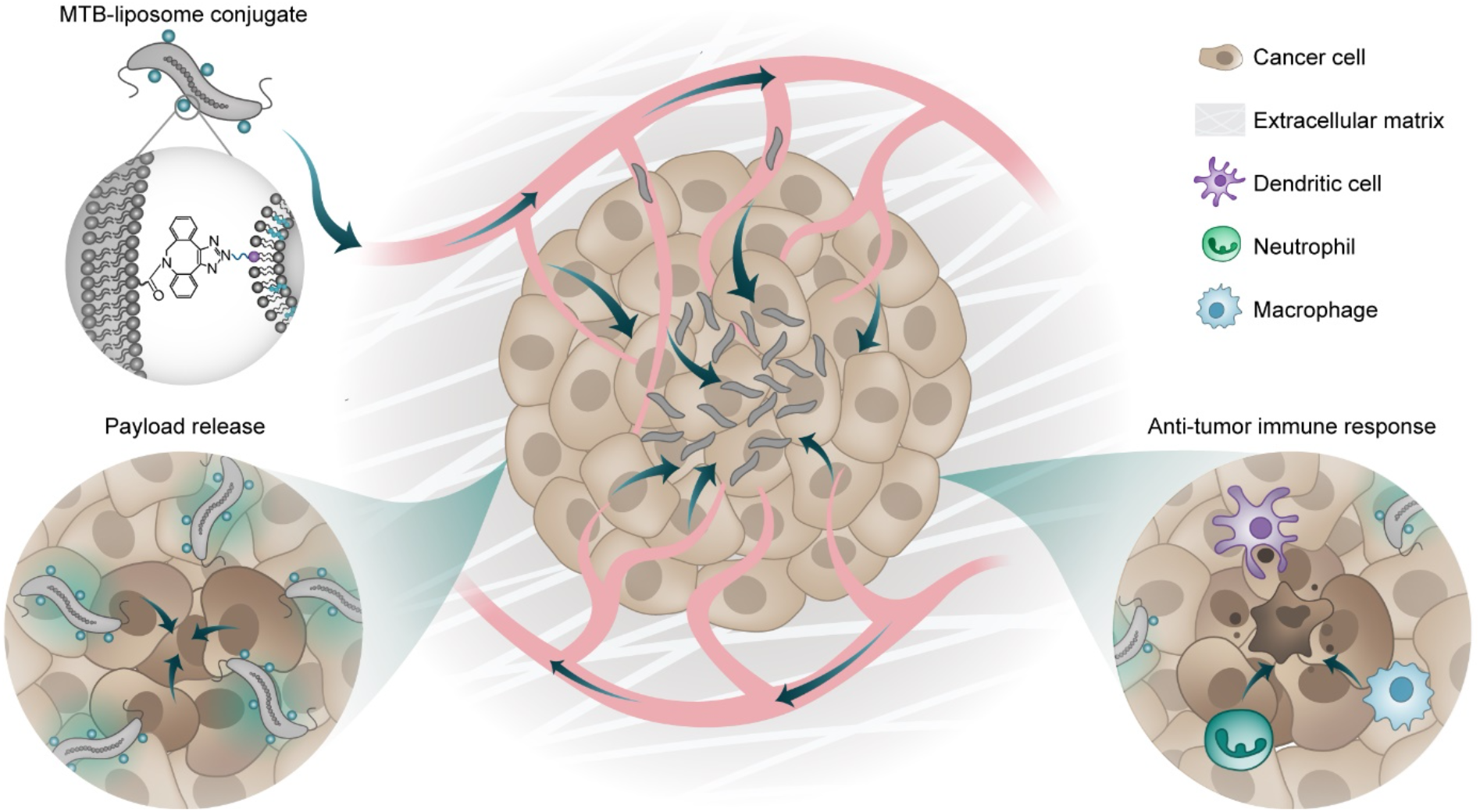
Conceptual overview of envisioned strategy using MTB-LP as living immunotherapeutic agents. Illustration of MTB biohybrid microrobots bearing drug-loaded liposomes conjugated to the cell membrane via a copper-free click reaction. Schematics depicting bacterial tumor colonization followed by payload release and immune cell recruitment.

**Figure Error!.**
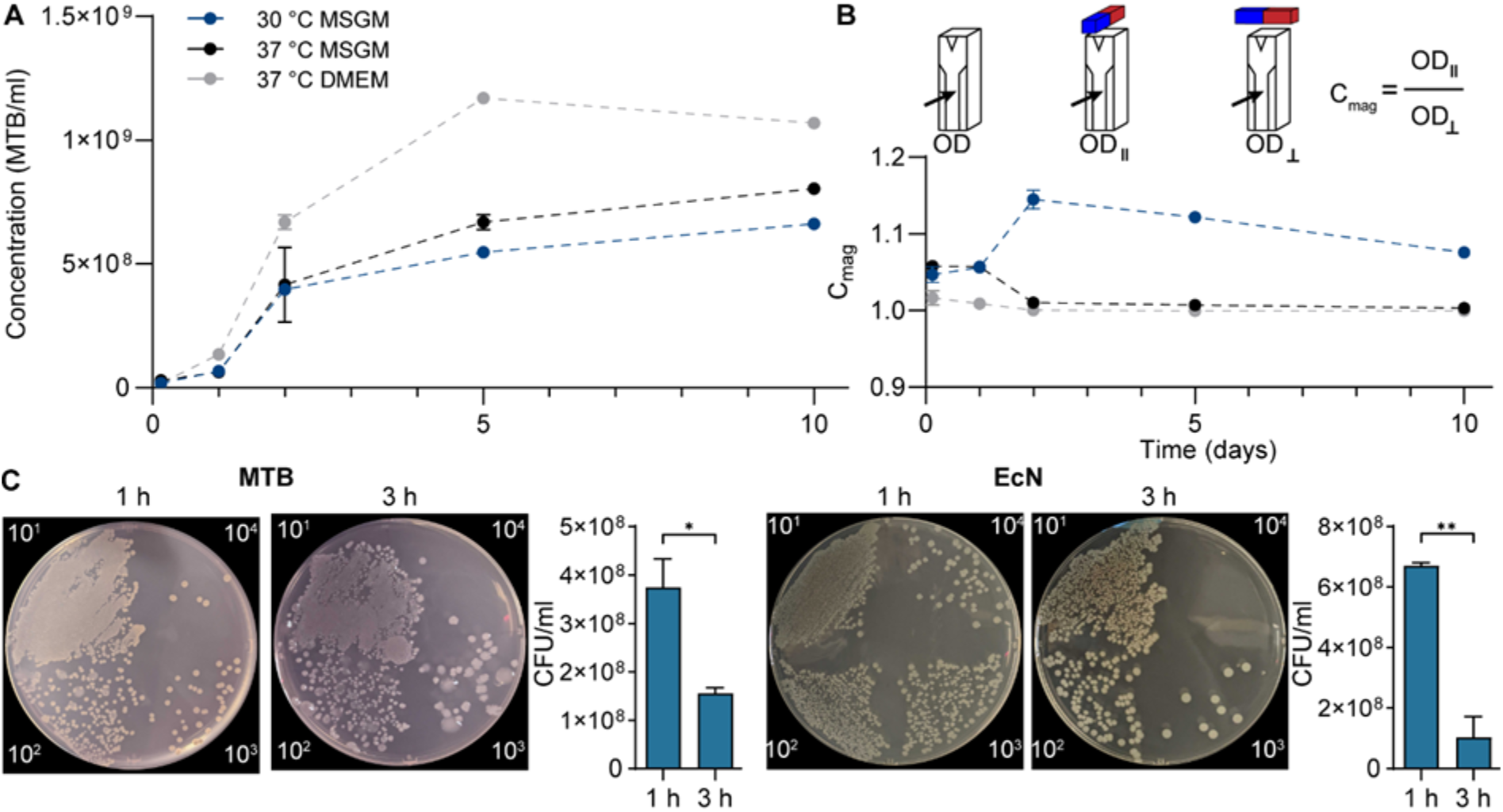
No text of specified style in document. Assessing MTB proliferation, magnetic responsiveness, and viability under varying conditions. (A) Bacterial proliferation under different culture conditions. OD was measured after 0.125, 1, 2, 5 and 10 days of incubation and used to compute bacterial concentrations (n = 3; mean ± SD). (B) Schematic illustrating OD and Cmag measurements (top). Magnetic responsiveness of bacterial populations, as indicated by Cmag values, cultured under different conditions over 10 days (bottom; n = 3; mean ± SD). (C) Representative images of MTB and EcN grown on agar plates following 1 or 3 h exposure to human whole blood. Samples were serially diluted the number of viable bacteria was quantified. (n ≥ 2 donors; mean ± SD; *P < 0.05 and **P < 0.01, Student’s t-test).

A unique characteristic of MTB is their ability to biomineralize iron oxide nanoparticles that render them magnetically responsive. C_mag_ values were used to assess the magnetic properties of bacteria cultured under different conditions (Figure 2B). C_mag_ is the ratio of light absorbance measurements when a magnet is placed parallel and then perpendicular to the light path, with values above 1 indicating the presence of magnetic material with geometric anisotropy in the suspension. MTB cultured under standard conditions (30 °C MSGM) consistently had C_mag_ values above 1 with a peak after 2 days in culture, indicating a magnetically responsive population. Samples cultured at 37 °C had C_mag_ values of approximately 1 by day 2, signifying that the bacterial suspensions were no longer magnetically responsive.

These findings suggest that while MTB have higher rates of proliferation at 37 °C in DMEM, biomineralization is impaired at this temperature. The MTB strain AMB-1 is typically cultured between 25 °C and 30 °C in iron rich media under microaerophilic conditions (*18*). Calugay *et al*. found that MTB proliferation increased and magnetosome formation decreased in low iron media (*19*), such as DMEM. Since magnetic cells biosynthesize highly organized magnetite, their growth rate is slower than that of non-magnetic cells. However, temperature appears to play a major role in the inhibition of magnetosome formation, as cultures at 37 °C in MSGM also had reduced magnetic responsiveness and higher rates of proliferation. The elevated temperature may induce stress and shift focus from the production of proteins for magnetosome synthesis to maintaining vital functions and preserving cell integrity (*20*). These findings support the use of MTB for bacteria-based cancer therapy since increased proliferation at 37 °C may assist in the formation of robust colonies in the tumor niche, a prerequisite for effective treatment.

To work towards the use of MTB for cancer therapy applications *in vivo*, a whole blood assay was performed to track bacterial viability over time (Figure 2C). Following intravenous administration, MTB would encounter various host bactericidal factors in blood. This study was used to gain insights into the amount of time that MTB might remain in circulation to assist in selecting an appropriate duration for magnetic actuation. Live EcN, a gram-negative strain frequently utilized for bacteria cancer therapy, was used for comparison. After 1 h incubation, high amounts of bacteria were present in all samples, while decreased concentrations were observed after 3 hours of incubation. The amount of viable MTB decreased 2.4-fold, while EcN levels dropped sharply by over 6.5-fold. Overall, these findings suggest that magnetic intervention should be implemented within the first 3 h when the MTB population is still magnetically responsive and viable bacteria may still be in circulation.

### Assessing cytokine expression from macrophages

Having established that MTB proliferate under physiological conditions and remain viable in whole human blood, responses from relevant immune cells were investigated. Macrophages play a central role in detecting and tackling bacterial infections. The inflammatory response of human macrophages to MTB was assessed by measuring the level of various cytokines produced by THP-1-derived macrophages co-cultured with bacteria. THP-1 is a human monocytic cell line extensively used as a model system to study macrophage-related physiological processes and functions (*21*).

*In vitro* differentiation of THP-1 monocytes is most often achieved using phorbol 12-myristate-13-acetate (PMA) to produce cells with similar phenotype, cell morphology, surface marker expression and cytokine production as primary human macrophages (*22, 23*). Stimulation with live EcN was used for comparison and stimulation with LPS was used as a positive control. To avoid bacterial overgrowth that occurs at longer co-culture durations, an intermediate period of 9 h was selected and sufficiently high levels of cytokine production were captured for all three conditions compared to the media only control (Figure S2). A human cytokine array was used to measure the levels of several cytokines (Figure S3) and cytokines with 4-fold higher expression than the control were plotted and compared (Figure 3).

**Figure 3:**
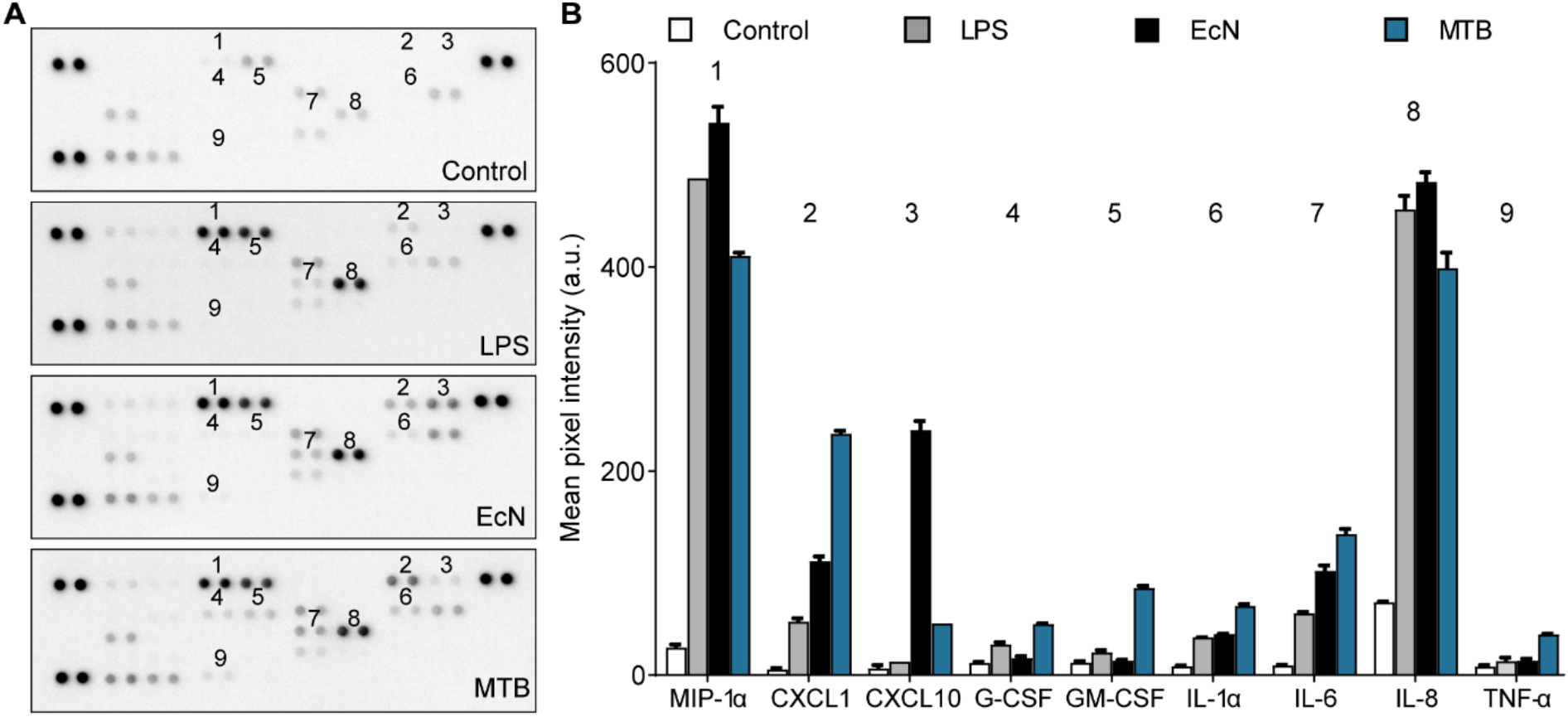
Assessing cytokine expression from THP-1 derived macrophages. **(A)** Human cytokine array performed on PMA differentiated THP-1 cells after 9 h of culture with media only, LPS, EcN or MTB. **(B)** Quantification of cytokine levels in THP-1 cell lysates using a human cytokine array. Cytokines with 4-fold higher expression than the control are plotted (mean ± SD of duplicates on array).

Stimulation with EcN and MTB produced similar levels of IL-8, MIP-1α, IL-1*α* and IL-6 expression. These cytokines are important mediators of proinflammatory responses and induce migration of polymorphonuclear cells, monocytes, DCs, natural killer cells, T cells, and platelets (*24–26*). IL-6, in particular, supports the growth of effector B cells and is antagonistic to regulatory T cells which prevent the development of effective antitumor immunity (*27, 28*).

MTB stimulation resulted in higher levels of TNF-α, CXCL1, G-CSF, and granulocyte-macrophage colony-stimulating factor (GM-CSF) expression than LPS and EcN. TNF-α has wide ranging effects, including causing necrotic regression of certain tumors (*29*). G-CSF and GM-CSF initiate the proliferation and differentiation of neutrophil progenitor cells, while CXCL1 induces neutrophil chemotaxis (*30–32*). In addition, GM-CSF stimulates the production of monocytes, which migrate into tissue and mature into macrophages and dendritic cells (*33*). Although MTB was not observed to induce substantial neutrophil migration in a Transwell assay (Figure S4), these findings suggest that higher levels of neutrophil recruitment are likely to occur.

EcN stimulation resulted in substantially higher levels of CXCL10 expression than MTB. CXCL10 is a chemoattractant for APCs, NK cells and T cells, and is associated with antitumor activity, although protumor effects have also been observed (*34*). Interestingly, it has been found to have direct antimicrobial functions, likely mediated through the cationic domain on the molecule causing disruption to the negatively charged bacterial cell wall (*35*). Thus, low levels of CXCL10 may be beneficial for bacteria-mediated therapies.

Overall, the proinflammatory cytokines expressed in response to stimulation with MTB were in line with those typically produced in response to other bacterial infections (*36*). Moreover, MTB stimulation generally did not result in substantially higher cytokine expression compared to EcN. While cytokine production is essential for pathogen clearance, excessive immune cell recruitment and cytokine release can lead to capillary leakage, dysregulation of immune responses and tissue damage (*37*). This characterization and establishment of a profile of human cytokine expression could assist future efforts to determine the safety and efficacy of MTB, and may also aid in identifying suitable candidates for effective combinational therapy.

### MTB promote cancer cell material uptake by moDCs

Dendritic cells are of interest in the context of bacteria-mediated therapy since they participate in responses to bacterial infections and also play a pivotal role in mounting an adaptive immune response against cancer. The uptake, processing, and subsequent presentation of cancer antigens by DCs to effector cells is a critical step in this process. Naïve DCs undergo maturation upon encountering various inflammatory mediators, after which the maturation marker CD83 is upregulated. Mature DCs have increased capacity to migrate towards lymphoid tissues and prime T cells (*38*). Since circulating DCs are scarce and difficult to isolate, many *in vitro* experiments are conducted using monocyte-derived DCs (moDCs) (*39*). MoDCs originate from monocytes that have been recruited to sites of injury or inflammation and differentiate into cells with a DC phenotype in response to cytokines in the surrounding environment (*38*) (Figure S5).

The uptake of cancer cell material by moDCs can be studied by staining both cell types and quantifying the percentage of double stained cells using flow cytometry (*40*). The ability of MTB to increase uptake was investigated using moDCs cultured with cancer cells in the presence of MTB. Two co-culture conditions were investigated (Figure 4A). The first condition, where MTB and CFSE-stained moDCs were placed in culture simultaneously with far-red-stained HCT116, was designed to represent a tumor microenvironment where DCs are already present. The second condition, where the cancer cells are co-cultured with MTB prior to the addition of moDCs was designed to represent a TME where moDCs are recruited after bacterial colonization. Since MTB was found to induce moDC maturation within 6 h (Figure S6) and mature moDCs have reduced phagocytic capacity (*38*), a duration of 4 h was selected for moDC co-culture in both conditions.

**Figure 4:**
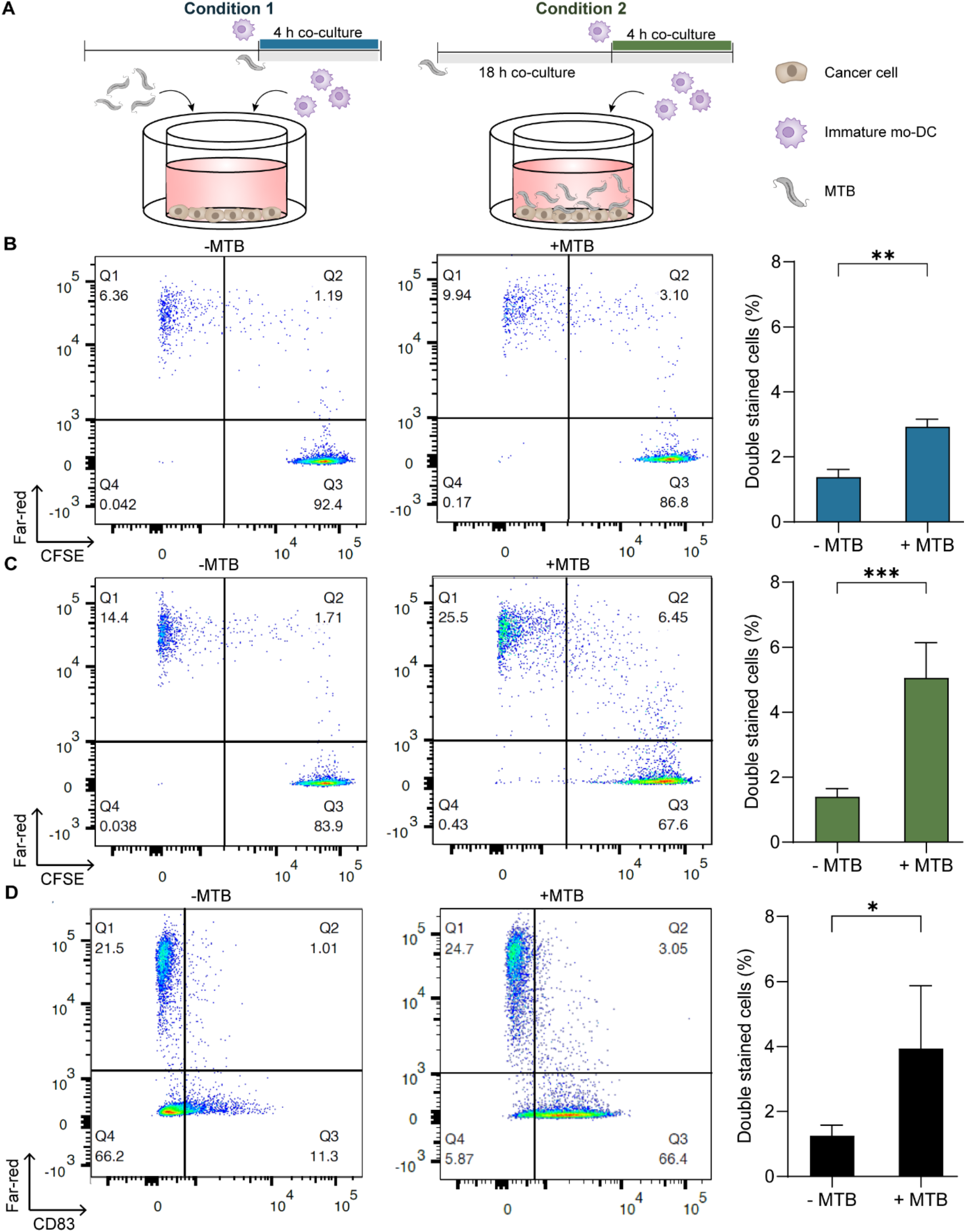
Cancer cell material uptake by primary human moDCs in co-culture with MTB. **(A)** Schematic illustrating two experimental conditions where stained moDCs and far-red stained HCT116 cancer cells were used to assess uptake of cell material in the presence of MTB. **(B)** Representative flow cytometry plots and quantification of cell material uptake for experimental condition 1, where MTB and CFSE-stained moDCs were added simultaneously (n = 2 donors; mean ± SD; **P < 0.01, Student’s t-test). **(C)** Flow cytometry analysis for condition 2 with prior MTB co-culture (n =3 donors; mean ± SD; ***P < 0.001, Student’s t-test). **(D)** Assessment of material uptake and CD83 expression for experimental condition 2 (n =3 donors; mean ± SD; *P < 0.05, Student’s t-test).

In the first condition, a 2-fold increase in the percentage of double stained cells was observed in the presence of MTB, indicating increased uptake of cancer cell material (Figure 4B). The effect was even more pronounced in the second condition, with an almost 4-fold increase in double-stained cells, suggesting that longer exposure of cancer cells to MTB promotes the uptake of cancer cell material (Figure 4C). Since the most substantial uptake was observed when cancer cells were cultured with MTB prior to co-culture with moDCs, this suggests that MTB had a direct effect on the cancer cells. In addition, CD83 expression was studied to assess whether maturation was induced in moDCs that phagocytosed cancer cell material. Mature DCs that have phagocytosed apoptotic cancer cells have been shown to mount highly efficient antitumor immunity (*41*). Approximately 90% of moDCs were CD83-positive when MTB was present during co-culture, while the majority of moDCs did not upregulate CD83 in the absence of MTB. Furthermore, 3-fold higher cancer cell material uptake by CD83+ moDCs was observed in cultures where MTB was present (Figure 4D). These results show that functional maturation of moDCs occurs, in addition to increased uptake of cancer cell material in the presence of MTB. To verify that active uptake of cancer cell material was responsible for the observed increase in double-stained cells, co-culture under the second experimental condition was also performed at 4 °C where the lower temperature inhibits phagocytosis (*42*) (Figure S7). At 4°C, decreased material uptake in the presence of MTB was observed, supporting the notion that MTB promotes active material uptake in moDCs.

### Covalent coupling of liposomes to MTB

Having established the suitability of the MTB strain AMB-1 for use in physiological environments, the use of MTB-LP as drug delivery agents was investigated. The MTB–LP complex combines the adaptability of liposomes with the functionality of magnetic-based platforms to produce a self-propelling, guidable agent for targeted drug delivery. The well-established thin film hydration method followed by sequential extrusion were used to fabricate monodisperse liposomes functionalised with azide groups (Figure S8). A copper-free click reaction was employed to covalently couple liposomes to MTB (Figure 5A). Click reactions offer high selectivity and this reaction was selected over copper-catalysed alternatives to preserve bacterial viability. In the first step of the reaction, MTB are functionalized with reactive dibenzocyclooctyne (DBCO) groups through an NHS reaction. Next, a copper-free click reaction between the DBCO-functionalized bacteria and azide-functionalized liposomes occurs, driven by the release of steric strain in the cycloalkyne molecule (*43*).

**Figure 5:**
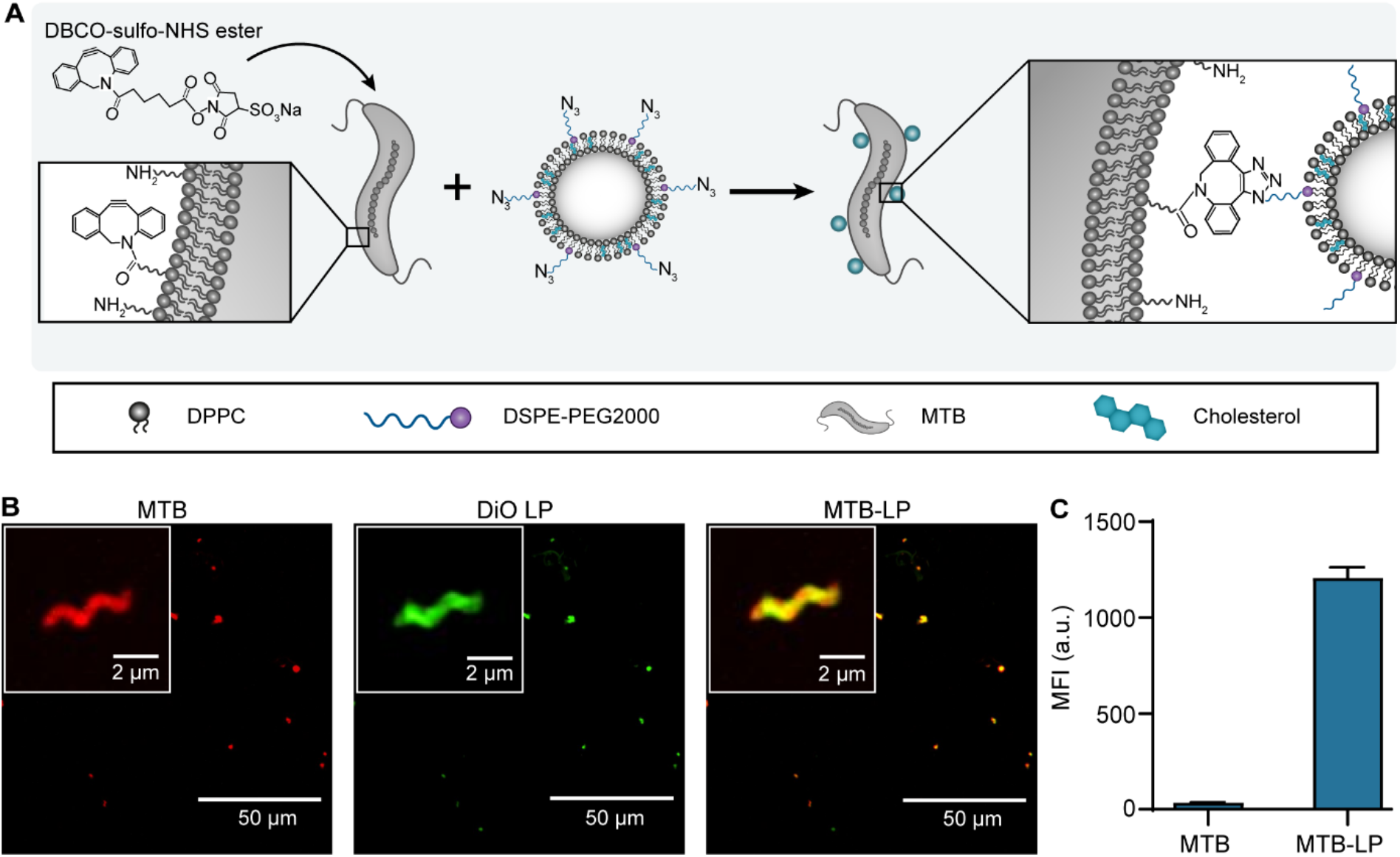
Covalent coupling to fabricate MTB-LP conjugates. (**A**) Illustration of copper-free click reaction to couple azide-functionalized liposomes to DBCO-functionalized MTB. (**B**) Confocal images of conjugates fabricated using a copper-free click reaction. MTB were stained with far-red and liposomes were tagged with DiO (DiO-LP; green). (**C**) Quantification of fluorescence intensity for MTB-LP with DiO-labelled liposomes (n = 3; mean ± SD; MFI = mean fluorescence intensity).

**Figure 6:**
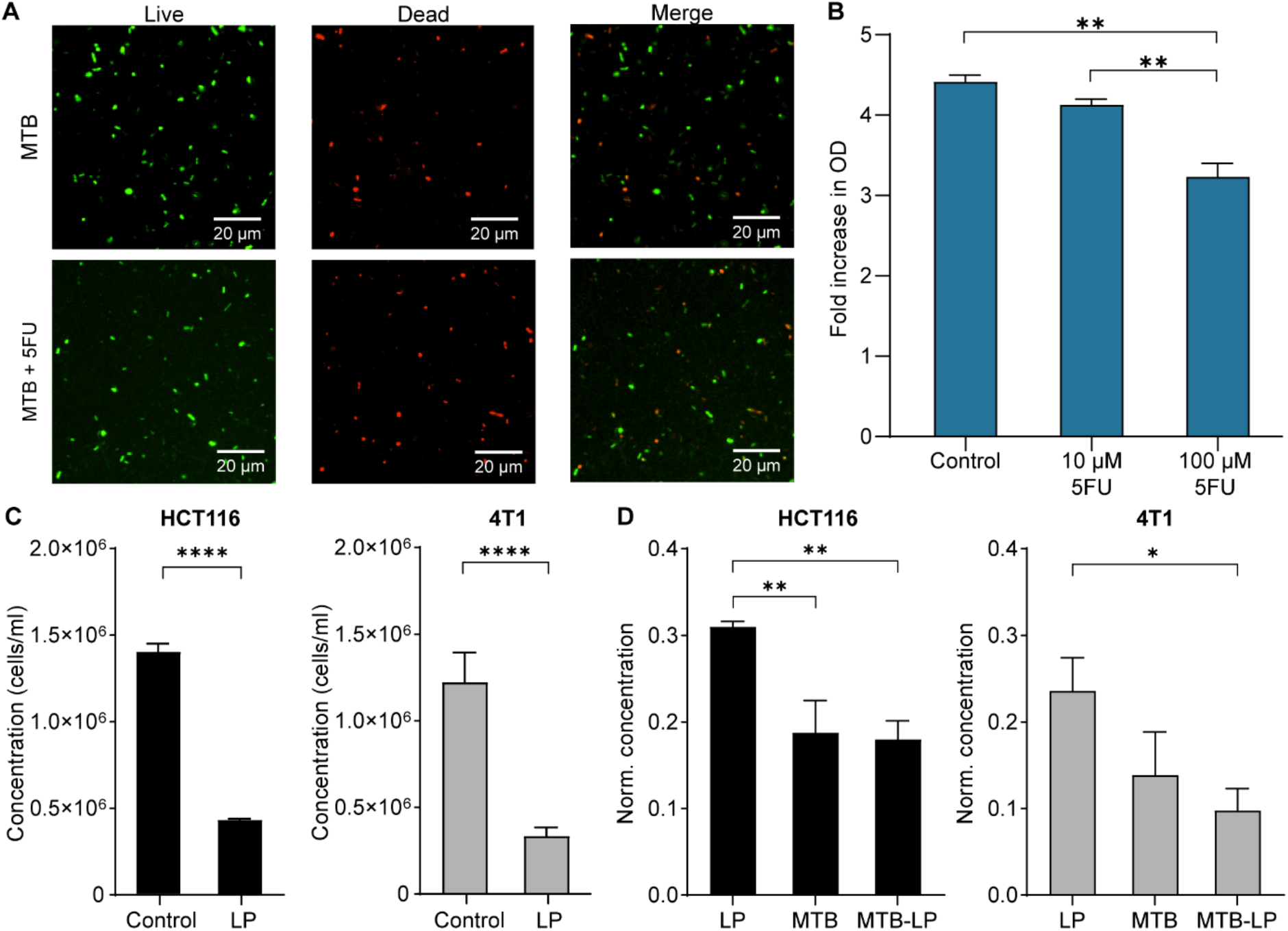
MTB-LP as delivery agents for 5-FU. **(A)** Confocal images of Syto9 and PI stained (dead) MTB following incubation with or without 10 μM 5-FU for 24 h in mammalian culture conditions. (**B**) Fold increase in OD_600_ measurements from 0 h to 24 h following incubation in media (control), 10 μM 5-FU or 100 μM 5-FU in mammalian culture conditions (n = 2; **P < 0.01 ANOVA). (**C**) Assessment of cell proliferation for HCT116 and 4T1 cells following 32 h incubation with or without 5-FU-loaded liposomes (LP) (n = 3; mean ± SD; ****P < 0.0001 ANOVA). (**D**) Cell proliferation normalized to untreated controls following 32 h incubation with LP, MTB or MTB-LP (n = 3; mean ± SD; *P < 0.05, **P < 0.01 ANOVA).

This reaction utilizes terminal amine groups present in phospholipids, LPS and various surface protein assemblies on the bacterial cell membrane (*44*). An estimate of the number of primary amine groups on the surface of MTB was determined using a reaction between primary amines with fluorescamine (Figure S9). The reaction produces a detectable fluorescent product and is extensively used to quantify amino acids and proteins (*45*). Since there was an insufficient amount of fluorescamine present to react with the excess amine groups in MTB suspensions at higher concentrations, these values represent an underestimation of the number of amines per bacterium. Therefore, lower MTB concentrations were used to provide an estimate of the number of amines per bacterium, which was found to be approximately 2.8 × 10^9^ amines/MTB.

Based on these estimates, conjugation of liposomes to MTB was performed. DiO, a green fluorescent, lipophilic carbocyanine dye, was incorporated into the lipid bilayer to enable imaging and quantification of MTB-liposome conjugation. Images showed co-localization of fluorescence signal from MTB and liposomes, and the resulting conjugates had a 35-fold higher fluorescence signal than MTB, confirming successful conjugation (Figure 3D).

### Complementary therapeutic effects of MTB-LP

Having developed an efficient method for generating MTB-LP, the possible complementary effect of MTB and a chemotherapeutic payload was explored. The widely used antimetabolite 5-FU was used as a model drug. Since 5-FU has been shown to have antibacterial activity by disrupting DNA metabolism in Gram-positive bacteria (*46, 47*), the effect of 5-FU on MTB was first investigated. To ensure that viability of Gram-negative MTB is not compromised by 5-FU, live-dead staining was performed. Images showed comparable amounts of dead cells in samples incubated with and without 5-FU for 24 hours (Figure 4A). Bacterial proliferation was also assessed (Figure 4B). Although MTB proliferation was reduced, particularly at higher concentrations, 5-FU did not completely inhibit bacterial proliferation or compromise viability, validating its suitability as a model drug for further studies.

It has been shown that 5-FU is most efficient when used in combination with other therapies (*48, 49*). Thus, the ability of MTB to enhance the therapeutic efficacy of the drug was investigated. As a basis for comparison, the effect of MTB-LP on proliferation was examined using cell lines with different p53 statuses. Although the antitumor activity of 5-FU is chiefly attributed to its ability to activate p53-dependent cell growth arrest and apoptosis, other mechanisms of action have been reported (*50*). Since 5-FU is the backbone of chemotherapeutic regimes for colorectal cancer, experiments were performed using the human colorectal cancer cell line HCT116, which has wildtype p53 (*51*). 4-T1 is a p53 deficient murine breast cancer cell line typically used as a syngeneic invasive cancer model (*52*). Using cell lines that require distinct pathways for 5-FU allows inferences to be made about the efficacy of the MTB-LP system for treating a wide range of cancer types.

The efficacy of the nanocarrier alone was first assessed and a significant decrease in proliferation for both cell lines was observed compared to untreated controls (Figure 4C). Next, the effect of MTB and MTB-LP were evaluated (Figure 4D). For both cell lines, co-culture with MTB resulted in a substantial reduction in cell proliferation, with a significant decrease observed in HCT116. Treatment with MTB-LP conjugates also resulted in a significant decrease in proliferation compared to liposomes only. For HCT116, the effect of MTB and MTB-LP were comparable, suggesting that most of the observed decrease in proliferation was a result of the presence of MTB. MTB have been shown to compete with cancer cells for essential nutrients like iron, thus acting as a self-replicating iron-chelator, and HCT116 cells are highly susceptible to treatment with iron chelators (*53–55*). In contrast, there was a notable decrease in proliferation following co-culture with MTB-LP compared to MTB alone for 4T1. Moreover, treatment with MTB-LP was shown to be significantly more effective than treatment with liposomes alone, suggesting that MTB-LP can improve the therapeutic efficacy of a delivered payload and could potentially be used for combination therapy.

### Conclusion

Although MTB are an atypical choice for bacteria-mediated therapy, the innate properties of MTB present a unique opportunity to facilitate targeted delivery and colonization of tumors. Their ability to migrate to low oxygen regions combined with their magnetic responsiveness can be leveraged for hybrid control strategies that merge magnetic manipulation with self-propulsion. Because research on the use of MTB for bacterial cancer therapy is still in its infancy, interactions between host immunity and MTB were yet to be studied. Stimulation of the immune system is an essential step for achieving tumor regression in bacterial cancer therapy. Thus, some key properties of human innate immune cells in response to stimulation with the MTB strain *M. magneticum* AMB-1 were examined in various co-culture assays. The findings of this work suggest a potential beneficial effect of MTB in mounting an immune-mediated antitumor response. Further characterization of the immunostimulatory properties of MTB is crucial for clinical use and will enable the safety and efficacy of this species to be determined. In addition, this work established the potential of the MTB-LP system as a versatile, complementary drug delivery platform. The results presented here provide evidence of a possible synergistic effect between MTB-LP and the delivered payload and suggest that using MTB-LP as delivery agents for chemotherapeutics might increase treatment efficacy. Overall, this study establishes the potential of the MTB-LP system as a versatile, combined drug delivery platform and lays the foundation for further characterization of the efficacy of MTB for immune-mediated cancer therapy.

## Materials and Methods

### Materials

HCT116 (human colorectal carcinoma, ATCC CCL-247), THP-1 (monocytic leukemia, ATCC TIB-202), *Magnetospirillum magneticum* (ATCC 700264), Wolfe’s Vitamin Solution and Wolfe’s Mineral Solution were purchased from American Type Culture Collection (Manassas, VA). The IVISbrite 4T1 Red F-luc Bioluminescent Tumour Cell line (Bioware Brite) was purchased from PerkinElmer (Waltham, MA). Fetal bovine serum (FBS) was purchased from BioWest (Nuaillé, France) and 4% formaldehyde (HistoFix 4%) was purchased from Carl Roth (Karlsruhe, Germany). Penicillin/Streptomycin (P/S) and transparent PET Transwell inserts were both purchased from Corning. Recombinant human anti-CD83 primary antibody, recombinant human anti-CD14 primary antibody, and goat anti-rabbit IgG (Alexa Fluor 488) secondary antibody were purchased from abcam (Cambridge, UK). Dulbecco’s Modified Eagle’s Medium (DMEM), McCoy’s 5A medium, Roswell Park Memorial Institute (RPMI) 1640 medium, lipopolysaccharide from E. coli (O111:B4) (LPS), 0.5 M EDTA, Pierce BCA Protein Assay Kit, CellTrace Far-Red and CFSE cell proliferation kits were acquired from Thermo Fisher Scientific (Waltham, MA). The human IL-1β antibody and goat IgG HRP-conjugated antibody were purchased from R&D Systems. Potassium phosphate, succinic acid, tartaric acid, sodium nitrate, ascorbic acid, sodium acetate, resazurin sodium salt, Luria-Bertani (LB) broth, LB agar, ferric quinate, agar, sodium hydroxide (NaOH), 3,3′-Dioctadecyloxacarbocyanine perchlorate (DiO), puromycin, 5-fluorouracil (5-FU), fluorescamine, Histopaque-1077, Histopaque-1119, recombinant human granulocyte-macrophage colony-stimulating factor (rhGM-CSF), recombinant human IL-4 (rhIL-4), N-Formyl-Met-Leu-Phe (fMLP), human serum albumin (HSA), sodium deoxycholate, IGEPAL CO-520, Tris(hydroxymethyl)aminomethane (Tris), sodium chloride (NaCl), sodium dodecyl sulfate (SDS), cholesterol, bovine serum albumin (BSA), phorbol 12-myristate-13-acetate (PMA), phosphate-buffered saline (PBS), were all acquired from Sigma-Aldrich (St. Louis, MO). Precision Plus Protein All Blue Standards, Laemmli sample buffer, 2-mercaptoethanol, Mini-PROTEAN TGX Stain-Free Precast Gels 4-20%, Immun-Blot PVDF, EveryBlot Blocking Buffer, luminol-enhancer solution and substrate peroxide solution were purchased from Bio-Rad (Hercules, CA). 1,2-dipalmitoyl-sn-glycero-3-phosphocholine (DPPC) and 1,2-distearoyl-sn-glycero-3-phosphoethanolamine-N-[azido(polyethylene glycol)-2000] (ammonium salt) (DSPE-PEG2000-azide) were purchased from Avanti Polar Lipids, Inc. (Alabaster, AL). Sulfo-DBCO-NHS ester was purchased from Broadpharm (San Diego, CA).

### Mammalian cell culture

HCT116 cells were cultured in modified McCoy’s 5A medium supplemented with 10% FBS and 1% P/S. 4T1 cells were cultured in RPMI-1640 supplemented with 10% FBS and 5 µg/mL puromycin. Cells were cultured at 37°C and 5% CO_2_ in a humidified atmosphere. One day before experiments, media was replaced with antibiotic-free media. Cell concentration was determined using a haemocytometer and the required number of cells was seeded for each experiment as specified.

### Bacteria culture

*M. magneticum* strain AMB-1 was cultured in revised magnetic spirillum growth medium (MSGM) which contained 5.0 mL of Wolfe’s Mineral Solution, 0.45 mL of 0.1 % resazurin, 0.68 g potassium phosphate, 0.37 g succinic acid, 0.37 g tartaric acid, 0.12 g sodium nitrate, 0.035 g ascorbic acid and 0.05 g sodium acetate per litre of distilled water. The final pH was adjusted to 6.9 with 1 M NaOH before autoclaving. Prior to use, Wolfe’s Vitamin Solution (1000x) and 10 mM ferric quinate (200x) were added to the media. MSGM agar plates (0.7% agar) were prepared by adding 10x MSGM to an autoclaved solution of agar in DI water. MTB was grown under microaerophilic conditions at 30 °C and suspensions grown for 5 to 7 days were used for all experiments. *E. coli* Nissle 1917 cultures were grown overnight in LB media at 37°C with agitation at 225 rpm or on LB agar plates (1.5% agar) at 37°C.

### MTB proliferation in mammalian culture conditions

MTB was grown in varying media, temperature, and oxygen condition conditions, as indicated in Table 1. Bacterial proliferation was assessed by performing optical density measurements at 600 nm (OD_600_) (Spark multimode microplate reader, Tecan). C_mag_ measurements were performed to quantify the magnetic responsiveness of the samples by placing a magnet parallel (OD_||_) and then perpendicular (OD_ꓕ_) to the light path. The C_mag_ value was calculated as OD_||_ /OD_ꓕ_.

**Table 1:** List of MTB different culture conditions.

| Media | Supplements | Temperature | Oxygen condition | Media change |
| --- | --- | --- | --- | --- |
| MSGM | 2 mM Wolfe's vitamin solution<br>10 mM ferric quinate | 30 °C | Microaerophilic | No |
| MSGM | 2 mM Wolfe's vitamin solution<br>10 mM ferric quinate | 37 °C | Microaerophilic | No |
| DMEM | 10% FBS | 37 °C | Microaerophilic | No |
| DMEM | 10% FBS, 1% P/S | 37 °C | Microaerophilic | No |
| DMEM | 10% FBS | 37 °C | Microaerophilic | Yes |
| DMEM | 10% FBS | 37 °C | Normoxia | No |

### In vitro whole blood bactericidal assays

Collection of human peripheral blood samples was approved by the Kantonale Ethikkommission Zürich (KEK-ZH-Nr. 2021-00413) and signed informed consent was obtained from each donor. MTB and EcN suspensions were centrifuged at 10 000 x g and 5 000 x g respectively and the pellets were resuspended in PBS at OD_600_ = 1. Next, 1 µl of either MTB or EcN was added to 100 µl of whole blood in a 96-well plate and incubated at 37 °C for 1 or 3 h. Each sample was serially diluted and plated on agar plates. Colony counting was performed after overnight incubation at 37 °C for EcN samples or after 5 to 7 days incubation at 30 °C for MTB samples.

### Cytokine expression in THP-1 macrophages

THP-1 monocytes were seeded at a density of 5 × 10^5^ cells/mL in 6-well plates and differentiated into macrophages by adding 100 ng/mL PMA into the culture for two days. Differentiated macrophages were then cultured in the absence of PMA for 24 h prior to stimulation with 10 ng/mL LPS, 2.5 × 10^7^ EcN or 2.5 × 10^7^ MTB for up to 24 h. Cells were lysed with RIPA buffer (pH 8.0) containing 50 mM Tris, 100 mM NaCl, 0.1 % SDS, 1 % IGEPAL CO-520, 0.5% sodium deoxycholate, and 1 mM EDTA. Lysates were loaded in pre-cast gels for gel electrophoresis and transferred to PVDF membranes. Membranes were incubated in EveryBlot blocking buffer for 1 h and subsequently incubated overnight with an IL-1β primary antibody. Membranes were washed, incubated with a goat IgG HRP-conjugated secondary antibody for 1 h, developed with ECL Substrate and captured with a Biomolecular Imager (Azure Biosystems). Cytokine production was assessed using a human cytokine array kit (Proteome Profiler, R&D Systems) according to manufacturer instructions.

### Investigation of neutrophil migration in a Transwell assay

Neutrophils were isolated by layering 20 mL of diluted blood on a double gradient of 10 mL Histopaque-1077 and 10 mL of Histopaque-1119. Tubes were centrifuged for 30 min at 700 × g (brake off). The neutrophil layer was collected and washed in PBS with centrifugation for 10 min at 280 × g (brake off). Isotonic lysis of RBCs was performed twice by adding 9 mL of sterile DI water to 1 mL of the neutrophil suspension. The tube was agitated for 20 s before 1 mL of 10x PBS was added to restore isotonicity and the suspension was washed with PBS. The final pellet was resuspended in 0.5% HSA in RPMI.

Transwell inserts (pore size = 3.0 µm; pore density = 6 ± 2 × 10^5^/cm^2^) were incubated at 37 °C for 1 h in wells containing 2.5 µg/mL fibronectin in PBS, which was isolated from human plasma. The fibronectin-coated inserts were washed twice with PBS and left to dry overnight. Bacteria suspensions at a concentration of 7 × 10^8^ bacteria/mL and a 0.01 mM fMLP solution were prepared in RPMI with 0.5% HSA. To each well of a 12-well plate, 600 µl of each suspension and a media only control were added in duplicates. The plate was left for 1 h to allow the bacteria to settle. The coated Transwell inserts were then placed in the wells and 200 µl of neutrophil suspension at a concentration of 2 × 10^6^ cells/mL were added to the apical compartment of each insert. The plate was placed in an incubator at 37 °C and 5% CO2 for 2 h. The suspension from the basolateral chamber was collected and adhered cells were detached using 0.25% Trypsin-EDTA. The number of migrated cells in each sample was determined using a hemocytometer.

### Differentiation and maturation of monocyte derived dendritic cells

Peripheral blood mononuclear cells (PBMCs) were isolated by layering 16 mL of diluted blood on top of 10 mL of Histopaque-1077. Tubes were for centrifuged for 30 min at 400 × g (brake off). The PBMC layer was collected, washed with PBS twice and centrifuged for 10 min at 280 × g (brake off). The PBMC pellet was resuspended in PBS supplemented with 2% FBS and 1 mM EDTA and monocyte isolation was performed using the EasySep^TM^ human monocyte isolation kit (Stemcell Technologies) according to manufacturer instructions. Isolated monocytes were cultured in RPMI supplemented with 10% FBS, 1% P/S, 25 ng/mL rhGM-CSF and 20 ng/mL rhIL-4 at a density of 5 × 10^5^ cells/mL in 12-well plates. Half of the media was replenished after 2 days. Validation of moDC differentiation and maturation was assessed by staining for CD14 and CD83 on isolated monocytes, differentiated immature moDCs cultured for 5 days, and mature moDCs stimulated for up to 24 h with MTB or EcN at a ratio of 10:1 relative to the number of seeded monocytes or 100 ng/mL LPS. Cells were blocked with 1% BSA in PBS for 1 h, followed by incubation for 45 min at 4 °C with one primary antibody at a final concentration of 1 µg/mL. Cells were fixed with 4% formaldehyde and washed with PBS prior to incubation with an Alexa Fluor 488-conjugated anti-rabbit IgG secondary antibody (diluted 1:250). Flow cytometry was performed with a 488 nm excitation laser and 530/30 filter and 20 000 events were recorded per tube (LSRFortessa, BD Biosciences). Data was analysed using FlowJo (version.10.4.2, Tree Star).

### Cancer cell material uptake by moDCs in co-culture with MTB

HCT116 cells were cultured for 20 h at a seeding density of 5 × 10^4^ cells/well and subsequently stained with CellTrace far-red. Uptake of cancer cell material by primary human moDCs was investigated under two different co-culture conditions. In the first condition, MTB at a ratio of 100:1 with respect to the number of seeded cancer cells was added simultaneously with stained moDCs to the wells. In the second condition, MTB at a ratio of 100:1 was co-cultured with cancer cells for 18 h at 37 °C prior to adding immature moDCs. Immature moDCs were harvested after 5 days in culture and stained with CellTrace CFSE according to manufacturer instructions. For both conditions, moDCs were added to the wells at a ratio of 2:1 with respect to the number of seeded cancer cells and the plates were incubated for 4 h at 4 °C or 37 °C.

MoDC maturation was assessed after co-culture under the second experimental condition with unstained immature moDCs and MTB at a ratio of 200:1. After co-culture, CD83 antibody staining was performed as previously described. Analysis was performed by flow cytometry with a 488 nm excitation laser and 530/30 filter and with a 640 nm excitation laser and 670/30 filter to detect CFSE-far red or CD83-far red stained cells. At least 20 000 events were recorded for each sample and analysis was performed using FlowJo.

### Liposome fabrication and characterization

Liposomes consisting of DPPC, cholesterol (25 mol %) and DSPE-PEG2000-azide (5 mol %), and DiO (0.5 mol %) were prepared from a total of 14 µmol of lipids using thin-film hydration. Lipids were dissolved in chloroform at the relevant molar ratios and dried to a thin film under nitrogen with further vacuum desiccation overnight at room temperature. For imaging, 0.1 mol% DiO was combined with the lipids. The lipid film was hydrated with 1 mL 5-FU solution (5 mg/mL) and placed in a water bath for 1 h at 50 °C with continuous stirring. Liposomes were downsized using sequential extrusion on a heating block (Avanti) at 50 °C. For extrusion, liposomes were passed 21-times each through a 400 nm followed by a 200 nm polycarbonate membrane (Whatman). Unencapsulated payload was removed by washing 3 times with ultracentrifugation at 16,000 ×g for 20 min at 4 °C. The average diameter, size distribution and zeta potential of the liposomes were determined by dynamic light scattering (DLS) (Litesizer 500, Anton Paar).

### Quantification of primary amines on MTB

The number of primary amine groups on the surface of MTB was evaluated using a reaction with fluorescamine. MTB suspensions were washed three times in PBS with centrifugation at 9,400 ×g for 10 min. For the reaction, 100 µL of 10mM fluorescamine was added dropwise to MTB suspensions with final concentrations between 1 × 10^7^ and 1 × 10^9^ MTB/mL. After incubation for 5 min at RT, fluorescence measurements were performed at 430/465 nm using a multimode microplate reader.

### MTB-liposome conjugation

MTB were stained prior to conjugation for imaging and quantification, otherwise unstained bacteria were used. MTB at a concentration of 1.5 × 10^8^ cells/mL in PBS were stained using the CellTrace Far-Red cell proliferation kit. A stock solution was prepared according to the manufacturers protocol and 2 µL was added to the MTB suspension. Cells were incubated at room temperature for 20 min, protected from light with gentle agitation. After incubation, 10 µL of 1% BSA was added to the cell suspension for 5 min to remove free dye. Cells were pelleted and resuspended in PBS.

For the copper-free click reaction, 1.2 µmol of 150 mM sulfo-DBCO-NHS ester was added dropwise to 1 × 10^8^ MTB in 200 µL PBS (pH 7.4) and incubated for 30 min at RT. MTB was washed three times by centrifugation at 5,000 ×g for 10 min. DBCO-functionalized MTB were then incubated with azide-functionalized liposomes at a ratio of 1.75:1 azides to amines. Conjugates were washed three times by centrifugation at 5,000 ×g for 10 min. Fluorescence measurements were performed at 430/465 nm using a multimode microplate reader.

### Assessment of the effect of 5-FU on MTB proliferation and viability

MTB suspensions at OD_600_ = 0.12 were added to a 12-well plate and 5-FU was added to the wells at a final concentration of either 10 or 100 µM. After incubation for 24 h at 37 °C, proliferation was assessed by evaluating the fold increase in OD_600_. MTB viability was assessed using the BacLight bacterial viability kit (Invitrogen) according to manufacturer instructions.

### Assessment of effect of MTB-LP on cancer cell proliferation

HCT116 and 4T1 cells were seeded at a density of 50’000 cells/well and incubated overnight at 37 °C and 5 % CO_2_. Cells were cultured for up to 48 h in media (control) or with 5-FU LP, MTB or MTB-LP at a cell to MTB ratio of 1:100 or an equivalent amount of liposomes. Supernatants were collected and cells were harvested. Cell proliferation was assessed by comparing concentrations of the cell suspensions which was determined using a hemocytometer.

## Supporting information

**Figure S1:**
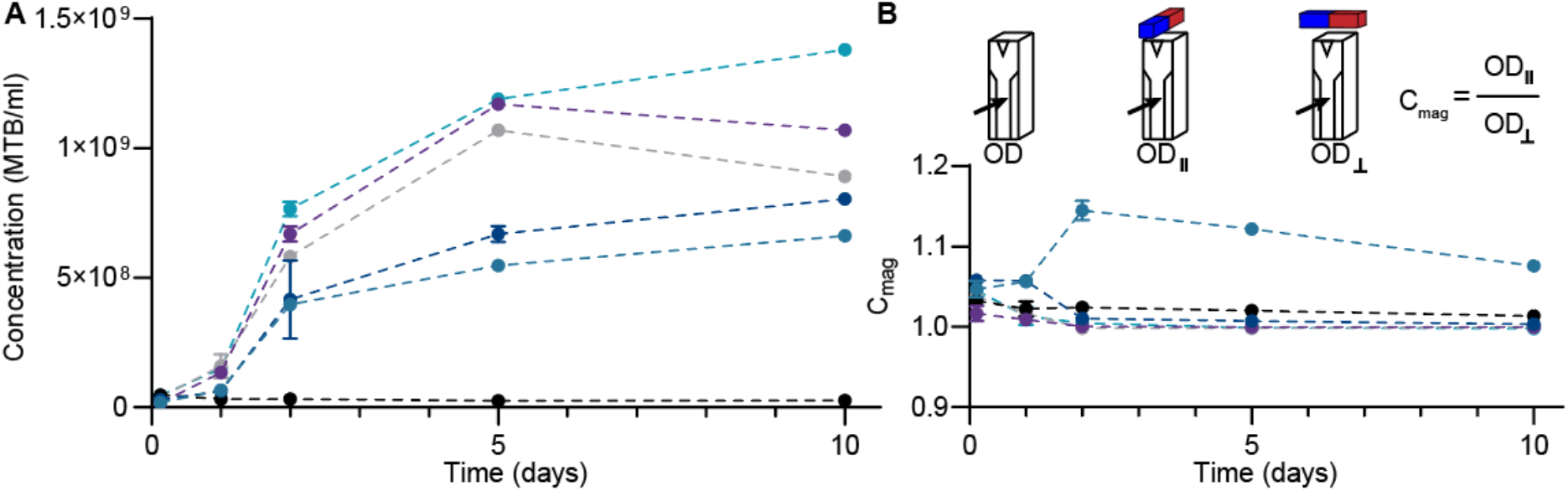
Assessing MTB proliferation and magnetic responsiveness under varying culture conditions. **(A)** Bacterial proliferation under different culture conditions. OD was measured after 0.125, 1, 2, 5 and 10 days of incubation and used to compute bacterial concentrations (n = 3; mean ± SD). **(B)** Schematic illustrating OD and Cmag measurements (top). Magnetic responsiveness of bacterial populations, as indicated by Cmag values, cultured under different conditions over 10 days (bottom; n = 3; mean ± SD).

**Figure S2:**
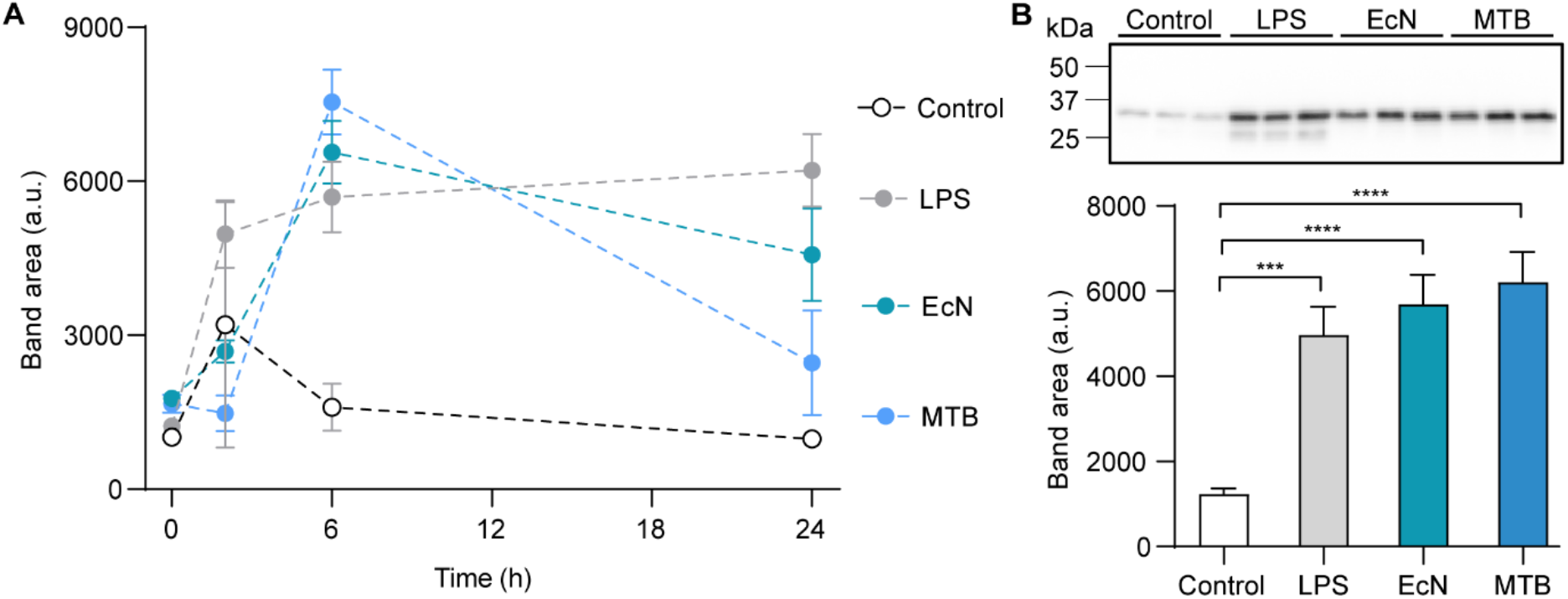
Assessing cytokine expression from THP-1 derived macrophages. (**A**) To determine the optimal co-culture duration, pro-IL-1β levels in THP-1 cell lysates following incubation with media only (control), LPS, EcN or MTB over 24 h was assessed using Western blotting (*n* = 3; mean ± SD). This cytokine is produced by macrophages in response to damage or pathogen recognition and is a key mediator of innate immune responses. (**B**) Pro-IL-1β levels assessed using Western blotting after 9 h of culture with media only, LPS, EcN or MTB (*n* = 3; mean ± SD).

**Figure S3:**
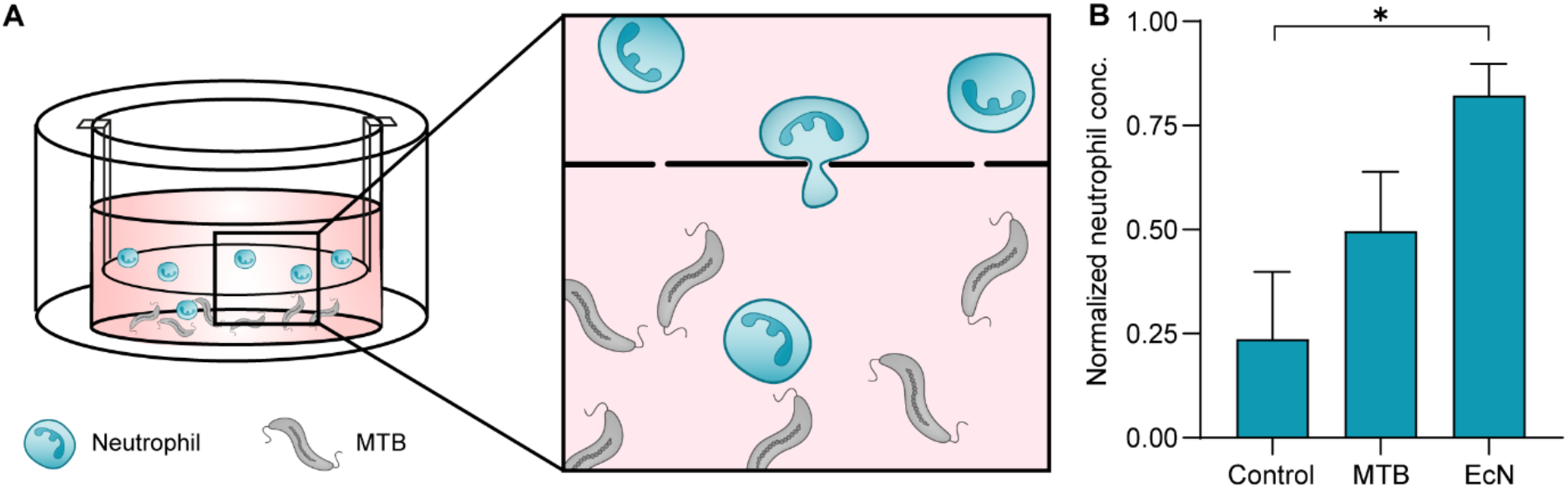
Cytokine expression in THP-1 derived macrophages. Quantification of cytokine levels in THP-1 cell lysates using a human cytokine array after 9 h of culture with media only, LPS, EcN or MTB (mean ± SD of duplicates on array).

**Figure S4:**
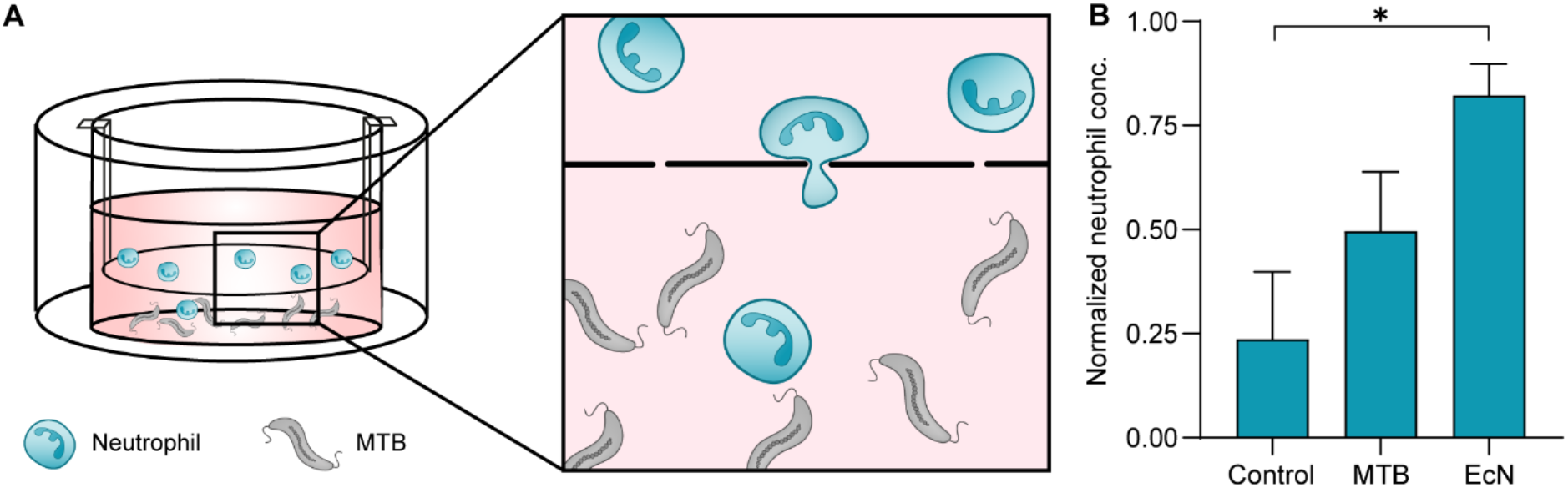
Assessing primary human neutrophil migration towards MTB. (**A**) Schematic representation of Transwell assay used to study neutrophil migration towards bacteria. (**B**) Comparison of normalized basolateral neutrophil concentrations following 2 h incubation in media only (control), MTB or EcN. Neutrophil concentration was normalized to concentrations resulting from directed migration along an fMLP gradient for each donor (n ≥ 2 donors; mean ± SD; *P < 0.05, ANOVA).

**Figure S5:**
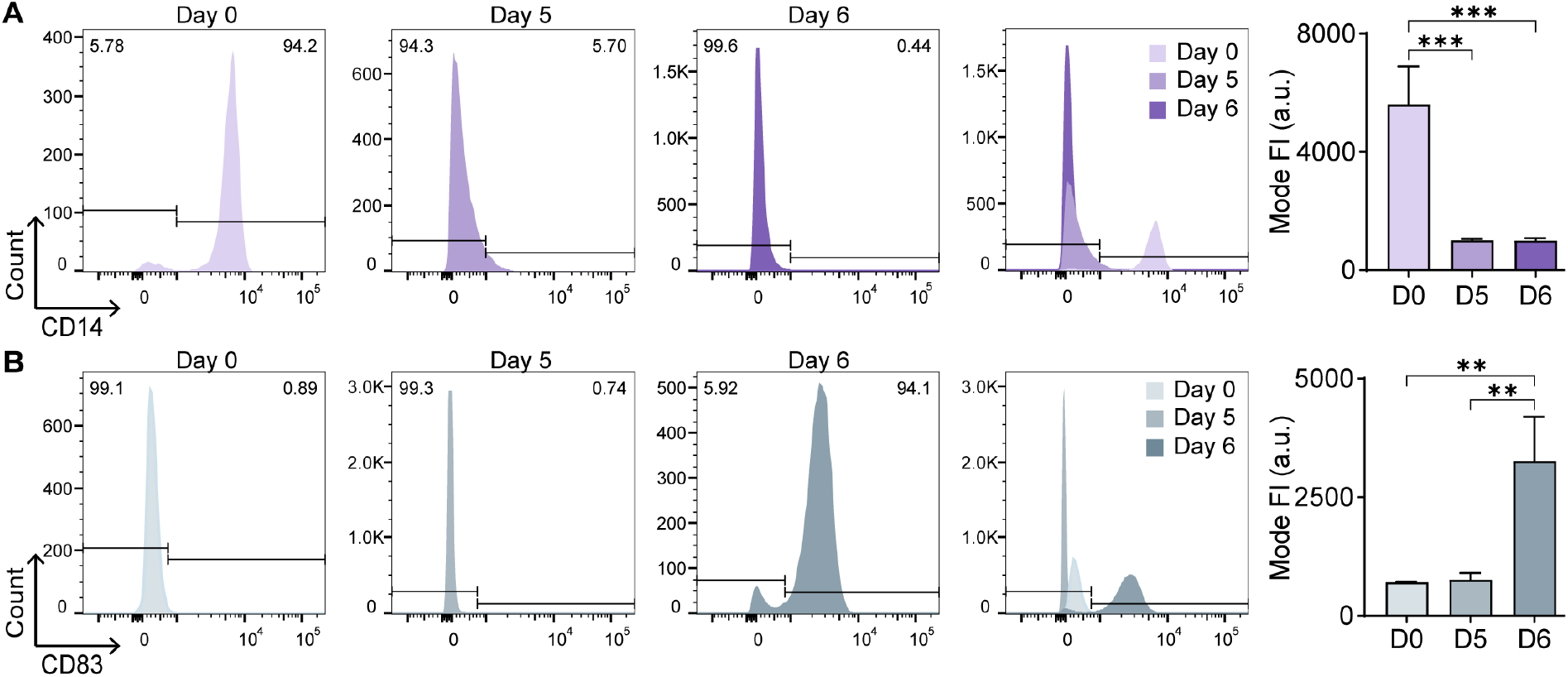
Validation of differentiation and maturation of primary human monocyte-derived DCs. **(A)** Representative flow cytometry histograms and quantification of CD14 expression in isolated monocytes (day 0), immature moDCs (day 5), and LPS-stimulated moDCs (day 6). FI= fluorescence intensity (*n* = 3 donors; mean ± SD; \*\*\**P* < 0.001, ANOVA). **(B)** CD83 expression after isolation (day 0) and during differentiation (day 5) and maturation (day 6) (*n* ≥ 2 donors; mean ± SD; \*\**P* < 0.01, ANOVA).

**Figure S6:**
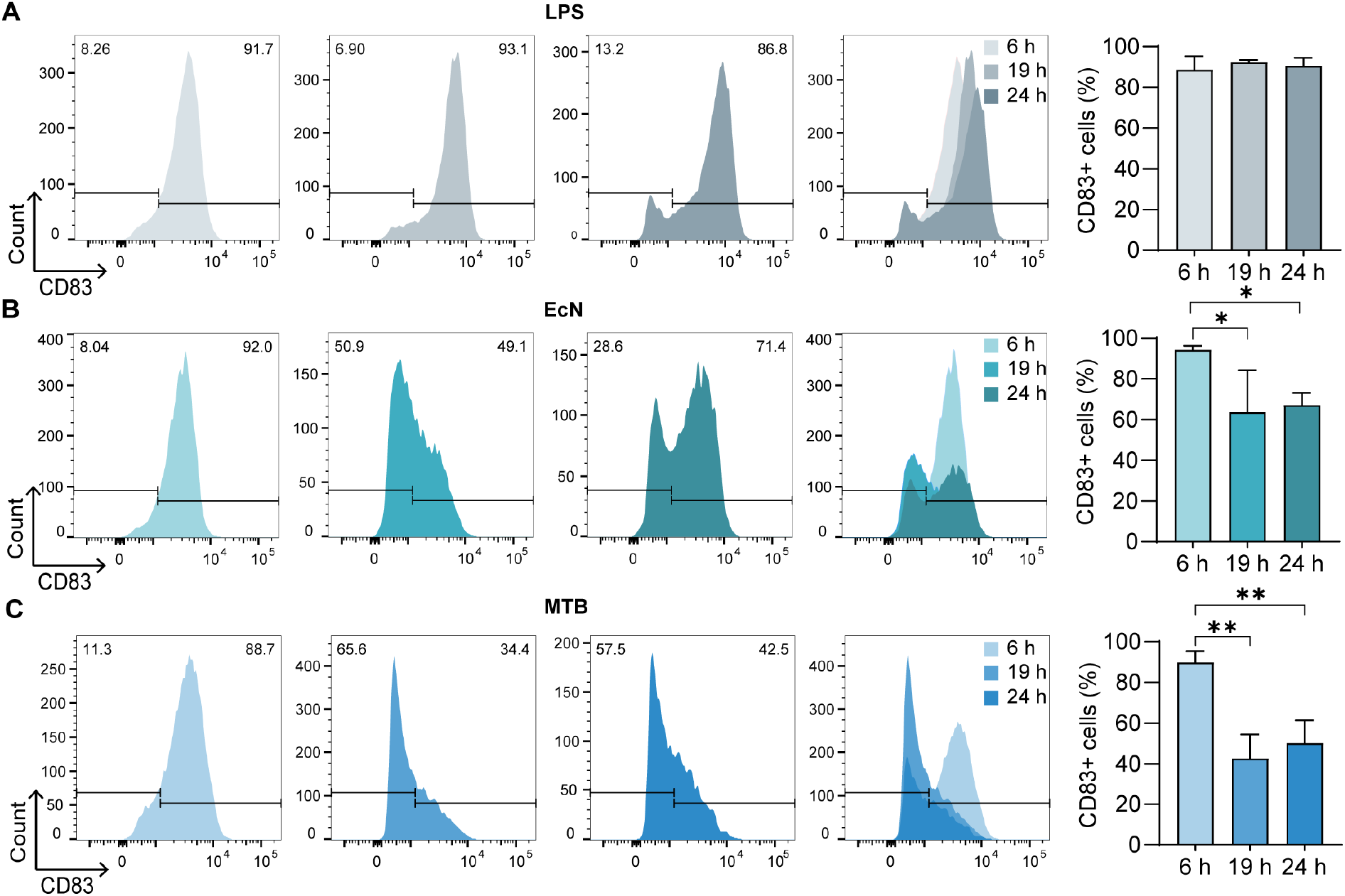
Assessment of primary human moDC maturation in the presence of MTB. (**A**) Representative flow cytometry histograms and quantification of the percentage of CD83+ moDCs following stimulation with LPS for 6, 19, and 24 h (*n* ≥ 2 donors; mean ± SD). (**B**) CD83 expression and quantification of CD83+ moDCs after co-culture with EcN for up to 24 h (*n* ≥ 2 donors; mean ± SD; \**P* < 0.05, ANOVA). (**C**) Assessment of expression and quantification of CD83+ moDCs after co-culture with MTB over 24 h (*n* ≥ 2 donors; mean ± SD; \*\**P* < 0.01, ANOVA).

**Figure S7:**
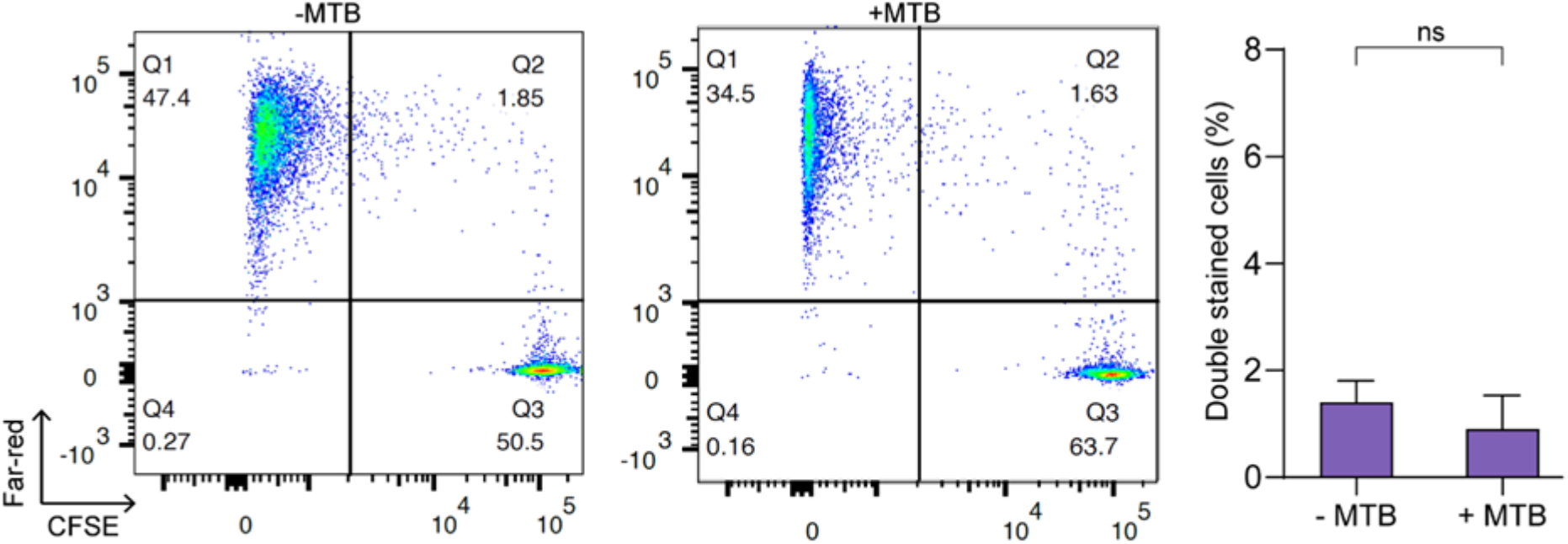
Validation of active uptake of cancer cell material by monocyte-derived DCs. Representative plots and quantification of material uptake for experimental condition 2 at 4 °C (n = 2 donors; mean ± SD; no significance, Student’s t-test).

**Figure S8:**
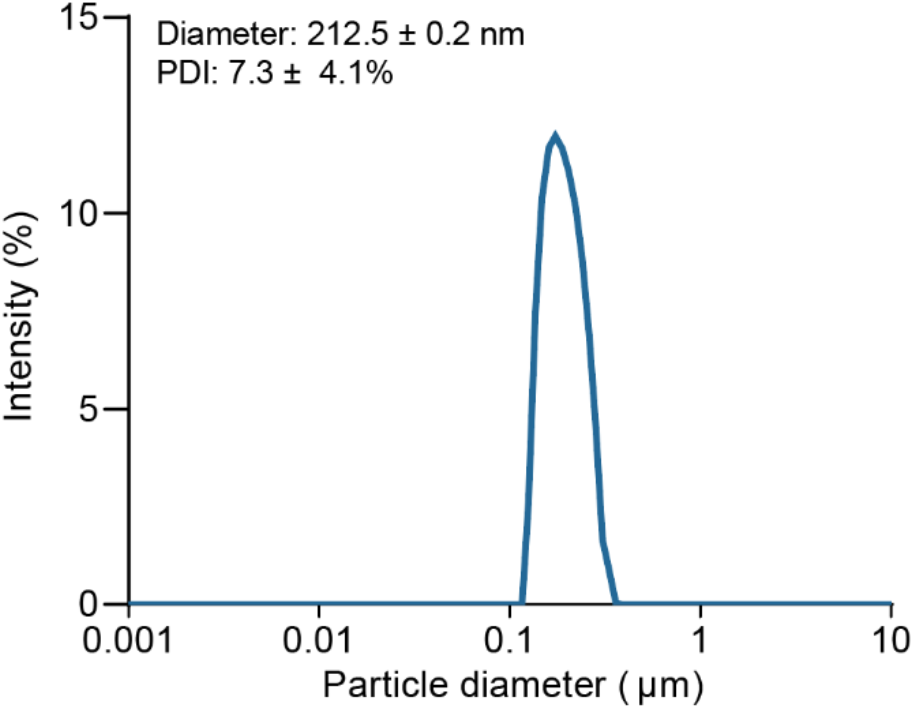
Characterization of fabricated liposomes. Average size distribution and polydispersity index (PDI) of liposomes functionalized with azide groups.

**Figure S9:**
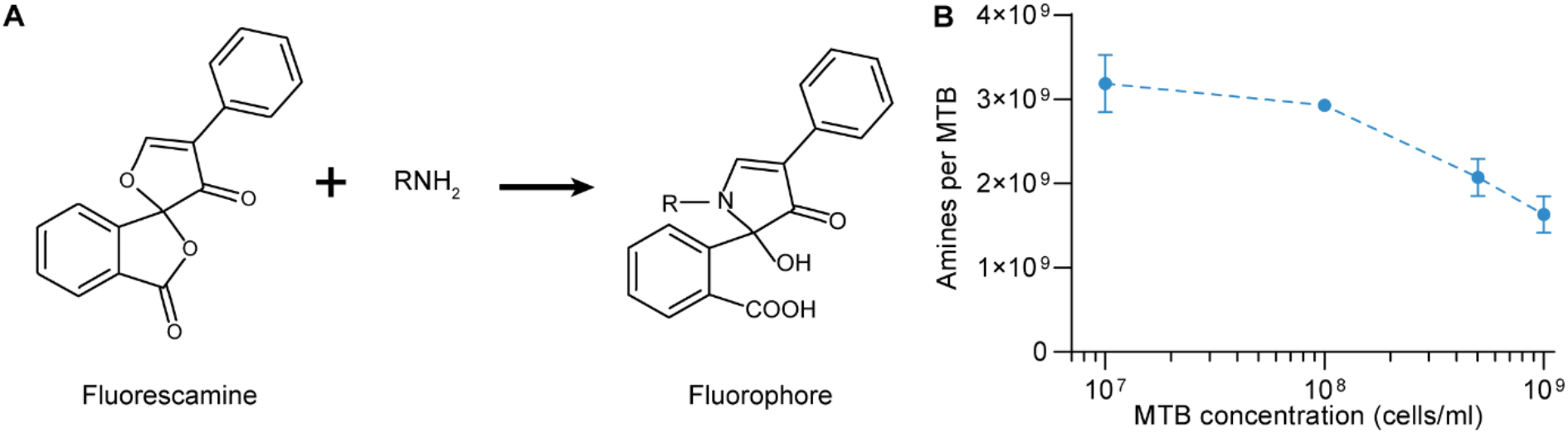
Quantification of amines on MTB cell membrane. (**A**) Reaction of fluorescamine with primary amines to produce a detectable fluorescent product. (**B**) Estimated number of primary amines per MTB at various bacterial concentrations (n = 2; mean ± SD).

